# The V617F mutation in JAK2 renders myeloid cells more sensitive to IL-6-mediated gp130 signaling

**DOI:** 10.1101/2025.10.10.681590

**Authors:** Henning Schurse, Marcelo A. S. de Toledo, Andrea Küster, Steffen Koschmieder, Madeline Caduc, Stefan Düsterhöft, Gerhard Müller-Newen

**Author notes:** Corresponding author: Prof. Dr. Gerhard Müller-Newen, Institute of Biochemistry and Molecular Biology, Uniklinik RWTH Aachen, Pauwelsstraße 30, 52074 Aachen, Germany. Shared senior authorship (Stefan Düsterhöft, Gerhard Müller-Newen).

## Abstract

The somatic V617F mutation in the pseudokinase domain of JAK2 (JAK2VF) causes various phenotypes of myeloproliferative neoplasms (MPN). By interacting with cytokine receptors such as those for erythropoietin (EPO) or thrombopoietin (TPO), JAK2VF induces ligand-independent dimerization and activation, leading to deregulated blood cell production, cytokine hypersensitivity, and inflammatory cytokine release. Interleukin-6 (IL-6), a key mediator of inflammatory symptoms in MPN, signals via homodimers of the gp130 receptor. We investigated whether JAK2VF alters gp130 dimerization and IL-6 sensitivity. Molecular dynamics simulations demonstrated that the JAK2VF pseudokinase domain forms more stable dimers than wild-type (WT) JAK2, potentially supporting gp130 tetramerization. In cell-based assays, IL-6 stimulation of JAK2VF+ cells induced stronger STAT3 activation than in JAK2-WT cells, reflecting enhanced IL-6 sensitivity. Moreover, JAK2VF expression elevated gp130 surface levels, dependent on the JAK2-binding motif in gp130. These findings indicate that JAK2VF promotes gp130 expression and dimerization, sensitizing mutant cells to IL-6. Thus, JAK2VF-driven amplification of IL-6/gp130 signaling may foster chronic inflammation and disease progression in MPN, representing a potential therapeutic target.

## 1. Introduction

Ph-negative myeloproliferative neoplasms (MPN) are a group of blood malignancies classified into three main subtypes: polycythemia vera (PV), essential thrombocythemia (ET), and primary myelofibrosis (PMF). These conditions are characterized by specific somatic mutations in the hematopoietic stem and progenitor cell compartment and their mature progeny, with the JAK2-V617F mutation (JAK2VF) being the most prevalent driver mutation occurring in 95% of PV and 55% of both ET and PMF cases [1]. The JAK2VF mutation affects the erythropoietin (EPO) receptor and the thrombopoietin (TPO) receptor, both of which are associated with the JAK2 tyrosine kinase. Under normal circumstances, upon stimulation with the respective cytokine, these receptors form homodimers resulting in activation of receptor-associated JAK2 and the downstream transcription factor STAT5, and in the case of TPO also STAT3 [2, 3]. The JAK2VF mutation renders these receptors constitutively active and hypersensitive to cytokine stimulation resulting in overproduction of erythrocytes and platelets [4, 5]. Among the numerous pro-inflammatory cytokines involved in MPN progression, IL-6 plays a prominent role in both CML [6] and Ph-negative MPN [7, 8]. Gp130 is the common receptor chain of the IL-6 cytokine family and gp130 signaling has been shown to critically depend on JAK1 [9] and JAK2 [10]. As gp130 also forms homodimers upon IL-6 stimulation, we investigated, whether the presence of JAK2VF makes cells hypersensitive to IL-6.

The family of JAK tyrosine kinases comprises four members: JAK1, JAK2, JAK3, and TYK2, each made of approximately 1,100-1,200 amino acids. The JAKs share a common structural architecture consisting of several distinct domains. The FERM (Four-point-one, Ezrin, Radixin, Moesin) domain at the N-terminus is followed by the SH2 (Src Homology 2) domain, which together form a tightly associated receptor-binding module. The pseudokinase domain (JH2), structurally similar to a tyrosine kinase domain but lacking enzymatic activity, plays a role in regulating the activity of the C-terminal kinase domain (JH1). This modular structure enables JAKs to bind to cytokine receptors and transduce signals via the JAK-STAT pathway [11]. The FERM-SH2 module forms an interface that binds the conserved Box1 and Box2 motifs in the membrane-proximal part of the intracellular cytokine receptor domain. Variations in the amino acid sequence of the Box1 and Box2 motifs facilitate binding specificity between cytokine receptors and JAKs [12, 13]. The JH1 domain contains the activation loop with two neighboring Tyr residues which become phosphorylated upon activation of the kinase (Y1007/Y1008 in JAK2) [14]. Mutational analysis focusing on the JH2 domain hinted at an autoinhibitory role, as JH2 deletion caused increased basal JAK activity and cytokine-independent STAT phosphorylation [15]. In 2005, multiple groups reported that the V617F mutation in JH2 of JAK2 induces aberrant JAK/STAT signaling and is the underlying cause for multiple forms of myeloproliferative neoplasms (MPN) [1, 16-18]. Structural analysis of the corresponding mutation in JAK1 (V657F) suggests that the replacement of valine by phenylalanine strengthens the interaction of a dimer interface in JH2 resulting in ligand-independent activation of the kinase [13]. In the present study, we analyze the consequence of the JAK2VF mutation for IL-6 signaling through the cytokine receptor gp130.

## 2. Results

### 2.1 *In silico* modeling predicts favored gp130-JAK2VF tetramerization

Understanding how individual Janus kinases (JAKs) interact with their cognate receptors remains a challenge, as key determinants are known, but comprehensive mechanistic insights are still lacking due to the limited number of experimentally solved structures. To address this gap, we used AlphaFold Multimer (AFM) to perform an unbiased AFM-based screen of either JAK1 or JAK2 against the cytosolic domains of homodimeric cytokine receptors including gp130 which forms a homodimer in the hexameric complex with IL-6 and its α-receptor (IL-6Ra) [19]. The AFM predictions mostly yielded high-confidence models suggesting favorable kinase-receptor dimer formation (Figure 1A). We validated the modeling approach by comparing one of our AFM predictions, the EPOR-JAK2 complex, to its experimentally determined structure (Figure 1B). This comparison showed a high degree of structural concordance. Moreover, the predicted dimer structures recapitulated known interaction motifs: in particular, the conserved Box1 and Box2 sequences of the receptor were correctly positioned at the interface with the JAK kinase. In the AFM models of JAK1-gp130 and JAK2-gp130, the Box1/Box2 region of the gp130 receptor were also sensibly docked to the JAK FERM-SH2 domains (Figure 1C), as expected from previous biochemical studies [20].

**Figure 1:**
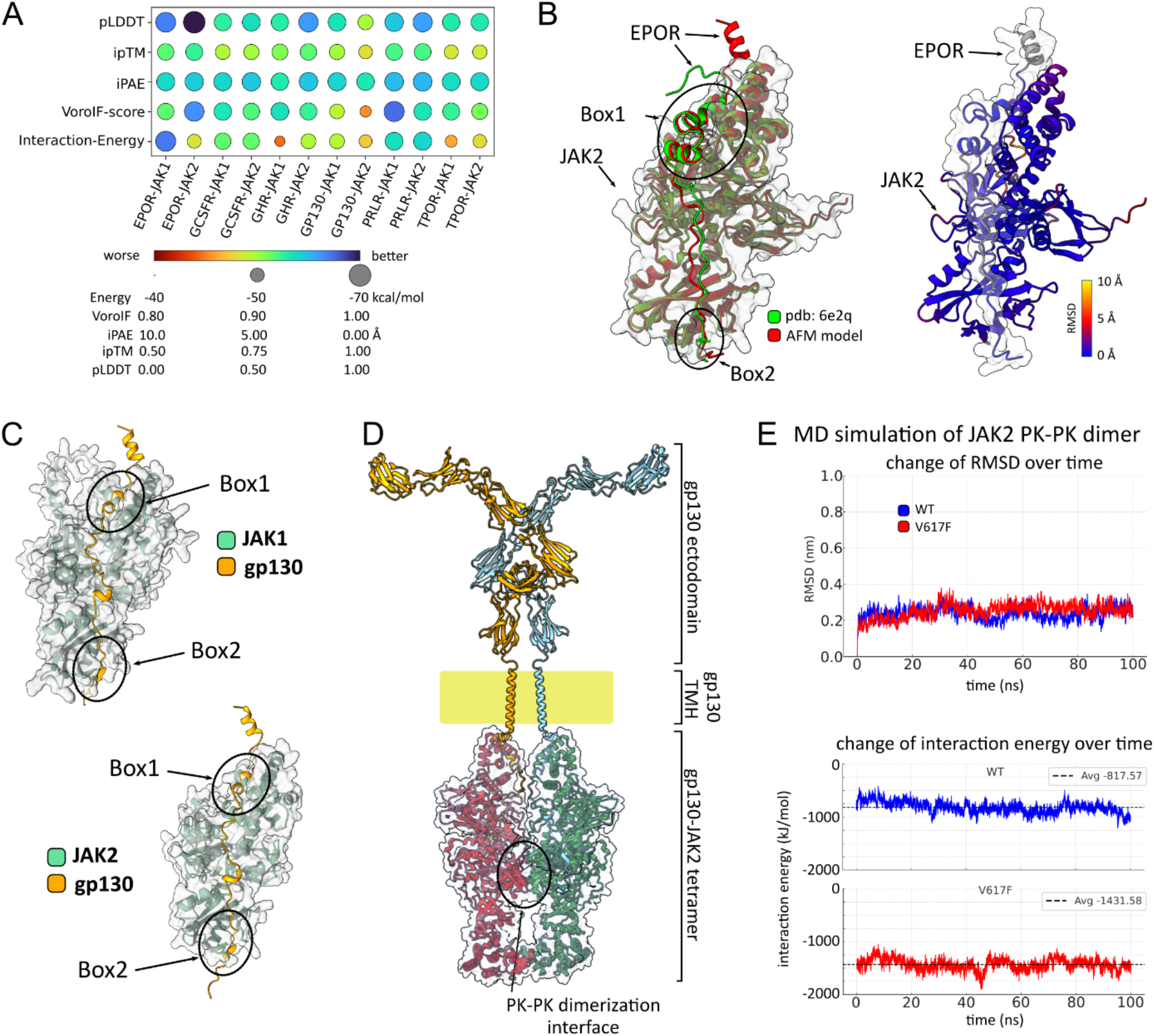
Kinase-receptor interaction modeling predicts favored gp130-JAK2VF tetramerization. **(A)** An unbiased screen for kinase-receptor dimerization was performed using AlphaFold Multimer (AFM)-based structural modeling. A bubble heatmap summarizes all analyzed dimer pairs with the evaluated metrics: AlphaFold Multimer scores (ipTM, pLDDT, iPAE), VoroIF-GNN scores, and FoldX calculated interaction energies. ipTM (interface predicted template modelling) score is an AFM interaction confidence score. pLDDT (predicted local distance difference test) score is a per residue structural confidence score. iPAE (interface predicted aligned error) estimates the positional error for each interface residue pair. VoroIF-score (weighted average predicted Contact Area Difference) is a confidence assessment score calculated by the AI-tool VoroIF-GNN [22]. Bubble heatmap shows differences by color gradient and circle size. **(B)** Validation of AFM predictions: (left) the AFM model of the JAK2-EPOR dimer (red) was compared with its experimentally determined structure (green, pdb: 6e2q). For this superposition the root mean square deviation (RMSD) values are 0.755 Å for 446 pruned atoms and 1.142 Å for all atoms. (Right) The AFM JAK2-EPOR dimer is colored by RMSD per residue. **(C)** AFM models of the JAK1-gp130 and JAK2-gp130 dimers are shown with the positioning of the Box1 and Box2 interfaces highlighted. **(D)** A composite AFM structure of the JAK2-gp130 tetramer with the PK-PK dimerization interface highlighted. **(E)** MD simulations of wild-type (WT) and JAK2VF PK-PK dimers: top graph shows that the RMSD reaches equilibrium within 10 ns. The bottom graph shows the interaction energies of the respective dimer. The interaction energies also remain stable over the runtime, with the JAK2VF dimer exhibiting a lower energy state (average interaction energy: 1431.58 kJ/mol) than WT (average interaction energy: 817.57 kJ/mol), indicating a more stable dimer interface in the JAK2VF dimer.

In addition to the internal AFM confidence scores, we applied complementary computational measures to further evaluate each modeled complex. Specifically, we used the FoldX force field to estimate binding interaction energies, and we applied VoroIF-GNN to evaluate the quality of the protein-protein interface. The results were consistent with known JAK biology: JAK2 was predicted to form stable complexes with the homodimeric receptors (Figure 1A). JAK1 interactions were predicted to be, in some cases, less favorable than those of JAK2, but still comparably stable. Importantly, other unknown determinants, not captured by the dimer structure modelling here, may further increase the preference of JAK2 over JAK1 for the homodimeric receptors. For gp130, the best ranking JAK partner was JAK1 followed by JAK2, consistent with literature [9, 10, 21].

Given the predicted binding of JAK2, we next asked how the oncogenic JAK2VF mutation affects JAK2 dimerization and JAK2-gp130 tetramerization (Figure 1D). We performed molecular dynamics (MD) simulations of the isolated JAK2 pseudokinase domain dimer, comparing the mutant (VF) and wild-type (WT) forms. The simulations showed that the interaction energy of the VF variant is much lower than that of the WT, demonstrating that the VF variant has an increased propensity for stable dimerization (Figure 1E).

In summary, our modeling results provide insights into how JAK family kinases may differentially engage their receptors. Moreover, the MD simulation shows that the JAK2VF mutation further stabilizes JAK2-JAK2 interactions, potentially facilitating the formation of higher order signaling assemblies such as gp130-JAK2 tetramers. This mechanism may increase cellular sensitivity to gp130-mediated cytokine signals.

### 2.2 Increased IL-6 response in 32D cells expressing JAK2VF

To investigate cytokine signaling via gp130 in the context of the JAK2VF mutation, we used 32D cells, a murine myeloblast-like cell line derived from bone marrow [23]. 32D cells were transduced with γ-retrovirus encoding either JAK2VF (JVF) or the empty transfer vector (EV). As 32D cells lack gp130 expression, 32DJVF or 32DEV (EV) cells were additionally transduced with γ-retrovirus encoding gp130. Flow cytometry analysis confirmed gp130 surface expression, which appeared to be stronger in 32DJVF cells compared to 32DEV cells (Figure 2A). As most cell types, 32D cells do not express the IL-6Ra. Therefore, these cells were stimulated with a combination of IL-6 and soluble (s) IL-6Ra to induce IL-6 trans-signaling [24]. Parental 32DJVF cells exhibited a basal induction of STAT3 phosphorylation due to constitutive JAK2 activity (Figure 2B, Figure S1). We did not observe an IL-6-mediated activation of STAT3 in any of the parental cell lines confirming the lack of gp130 expression. Importantly, 32DJVFgp130 cells showed a more pronounced STAT3 activation after IL-6/sIL-6Ra stimulation compared to 32DEVgp130 cells.

**Figure 2:**
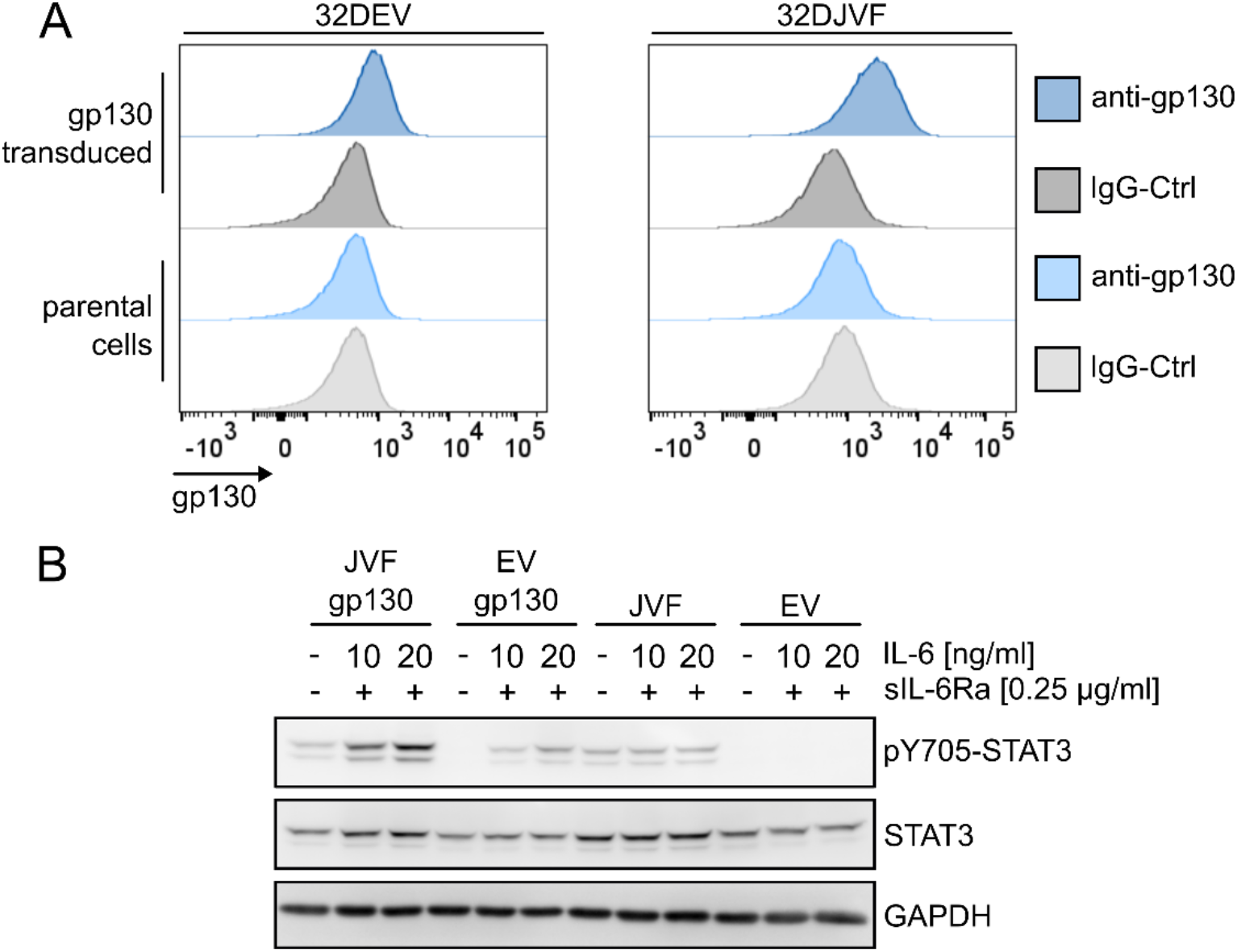
Increased IL-6 response in 32D cells upon expression of JAK2VF. 2DJAK2VF (JVF) and 32DEmpty-Vector (EV) cells were transduced with γ-retrovirus encoding human gp130. **(A)** Flow cytometry analysis of parental and gp130-transduced 32DEV or 32DJVF cells for gp130. Staining with an appropriate IgG antibody served as a control. **(B)** Transduced and untransduced cells were stimulated (10 or 20 ng/ml IL-6, 0.25 µg/ml sIL-6-Ra, 30’ at 37 °C) and cell lysates were prepared for SDS-PAGE and Western-blot analysis. The membrane was probed with a pY705-STAT3 antibody and detection of STAT3 or GAPDH served as controls.

### 2.3 Increased IL-6 response in JAK2VF+ cell is in-dependent of JAK1, but dependent on JAK2VF activity

Our interaction analysis of gp130 with JAK1 and JAK2 predicted that gp130 interacts more prominently with JAK1 than with JAK2 in agreement with previous experimental analysis of the gp130-JAK interactions [25, 26]. Functional analysis by gene targeting in mice revealed that JAK1 is the most relevant kinase for IL-6 signaling downstream of gp130 [21]. To specifically investigate the gp130-JAK2VF axis, we generated JAK1-KO 32DJVFgp130 and 32DEVgp130 cells by CRISPR/Cas9-mediated genome editing.

JAK1-KO was verified by Sanger sequencing (Figure S2B) as well as Western-blot analysis (Figure 3A) and Upon IL-6/sIL-6Ra stimulation, we again observed a stronger induction of pY705-STAT3 in 32DJVFgp130 cells compared to 32DEVgp130 cells (Figure 3A, Figure S3A). Stronger IL-6-mediated phosphorylation of STAT3 was also observed in 32DJVFgp130_CRISPR_ cells demonstrating that phosphorylation was JAK1-independent. Furthermore, 32DEVgp130_CRISPR_ cells responded to IL-6 stimulation with STAT3 activation as well, most probably mediated through endogenous JAK2. These findings underline that IL-6 can transduce signals in the absence of JAK1 and increased IL-6 sensitivity of 32DJVFgp130 cells is independent of JAK1.

**Figure 3:**
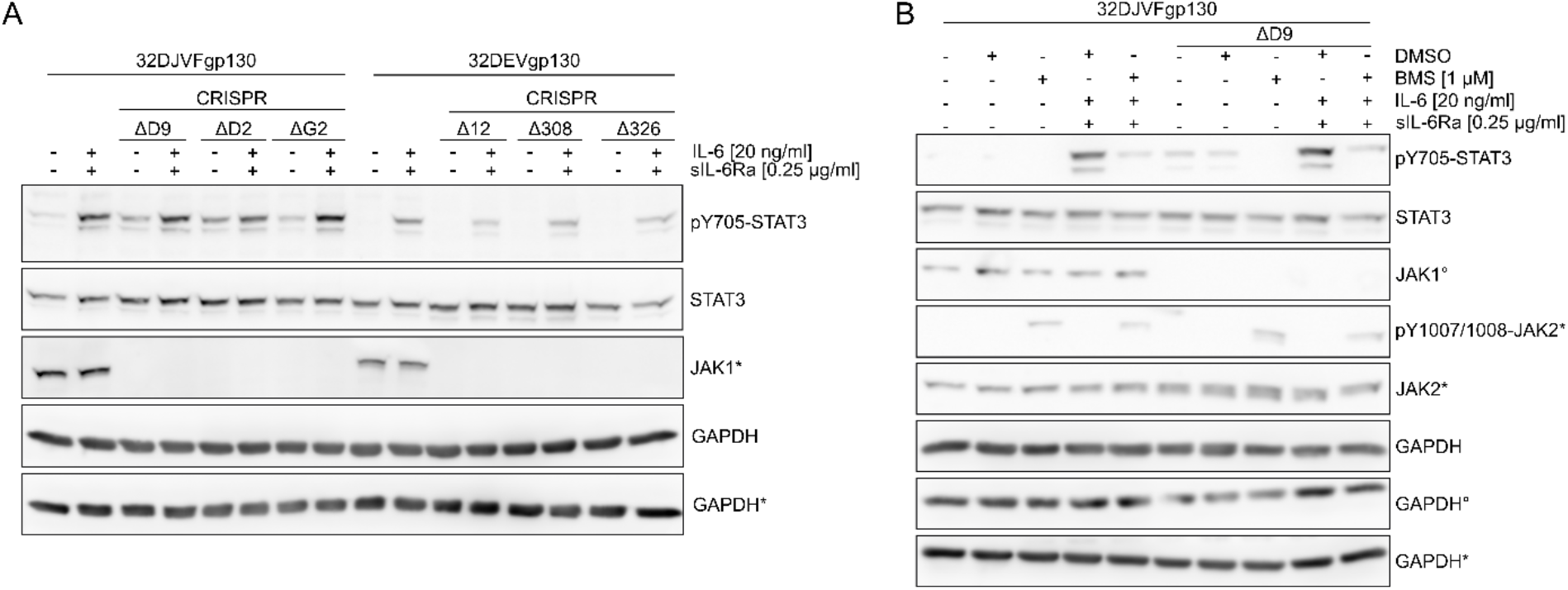
Increased IL-6-mediated STAT3 activation in 32DJVFgp130 cells is independent of JAK1 but sensitive to inhibition of JAK2. JAK1 knockout cells were generated via CRISPR/Cas9-mediated genome editing with subsequent amplification of single-cell clones (Δ). **(A)** Cell lysates were prepared from unstimulated and stimulated (20 ng/ml IL-6 and 0.25 µg/ml sIL-6Ra for 30’ at 37 °C) cells. SDS-PAGE and Western-blot analysis were performed for p705-STAT3 and JAK1. Detection of STAT3 and GAPDH served as controls. **(B)** Cells were treated with BMS for 16h prior to stimulation. Cell lysates were prepared for SDS-PAGE and Western-blot analysis of pY705-STAT3, pY1007/1008-JAK2 and JAK2. Detection of JAK2, STAT3 and GAPDH served as controls. Different immunoblots were used to detect STAT3 and JAKs. Corresponding loading controls are marked by * or °.

Next, we treated 32DJVFgp130 and 32DJVFgp130_CRISPR_ cells with the JAK2-specific inhibitor BMS-911543 (BMS) prior to IL-6/sIL-6Ra stimulation. As BMS is a type I inhibitor, it traps JAK2 in its active state causing an accumulation of pY1007/1008-JAK2 which is also observed with other type I inhibitors like ruxolitinib [27]. Importantly, pre-treatment with BMS prevented IL-6-mediated STAT3 induction in 32DJVFgp130 and 32DJVFgp130_CRISPR_ cells (Figure 3B, Figure S3B), indicating that IL-6 signaling in the absence of JAK1 is JAK2-dependent.

### 2.4 The JAK interaction motif in gp130 is required for increased gp130 surface expression and higher IL-6 sensitivity of JAK2VF+ cells

Both JAK2VF and CALR influence the surface expression of cytokine receptors such as TPOR [28, 29]. Therefore, we quantified the mean fluorescence intensity (MFI) for gp130 in our model cell lines, using flow cytometry. As expected, gp130-transduced cells exhibited a strongly elevated gp130 MFI compared to parental cells. Importantly, JAK2VF expression increased the gp130 MIF further, compared to EV controls (Figure S4A).

Next, we aimed to investigate whether increased gp130 surface expression required the interaction between gp130 and JAK2VF. To this end, 32DJVF and 32DEV cells were transduced with variants of gp130, either lacking the JAK interaction motif (ΔBox1) which abrogates JAK-binding, or carrying a mutation in the Box1 motif (W652A) which does not affect JAK interaction but downstream signaling [20]. Again, gp130-WT was significantly higher expressed at the cell surface in JAK2V617^+^ cells compared to EV cells (Figure 4A). However, surface expression of gp130-ΔBox1 was significantly lower compared to gp130-WT in JAK2VF^+^ cells. The surface expression of gp130-W652A was higher than gp130-ΔBox1 but lower than gp130-WT. To control for gp130 mRNA expression, we performed RT-qPCR and found similar gp130 transcript levels in all cell lines (Figure S4B).

**Figure 4:**
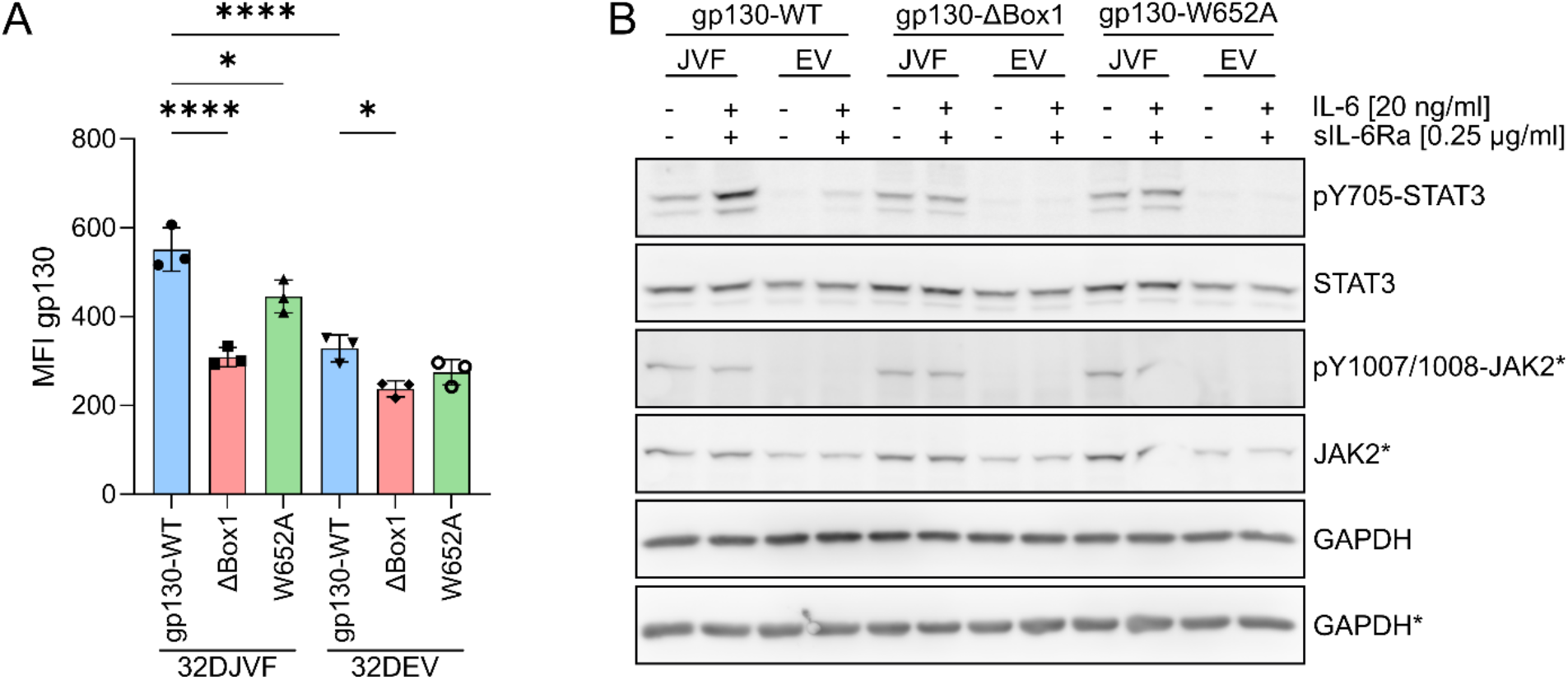
The JAK interaction motif in gp130 is necessary for increased gp130 surface expression and elevated IL-6 response. 32DJVF and 32DEV cells were transduced with gp130-WT, gp130-ΔBox1 or gp130-W652A. **(A)** Surface expression of gp130 in transduced cells was analyzed via flow cytometry. The geometric mean fluorescence intensity (MFI) of gp130 was calculated for all analyzed cell lines. Untransduced cells served as a control. **(B)** Indicated cell lines were stimulated with 20 ng/ml IL-6 and 0.25 µg/ml sIL-6Ra for 30’ at 37 °C. Cell lysates of unstimulated and stimulated cells were prepared for SDS-PAGE and Western-blot analysis of pY705-STAT3 and pY1007/1008-JAK2. Detection of STAT3, JAK2 and GAPDH served as controls. Different immunoblots were used to detect STAT3 and JAK2. Corresponding loading controls are marked by *. Data was analyzed via FlowJo^TM^ v10 as well as GraphPad Prism v10. P-values were calculated via one-way ANOVA with Tukey multiple comparison test (p<0.05*, p<0.0001****). Corresponding loading controls are marked by *.

Cells expressing the different gp130 variants were stimulated with IL-6/sIL-6Ra to investigate STAT3 activation (Figure 4B and Figure S5). Again, 32DJVFgp130 demonstrated an increased induction of pY705-STAT3. No stimulation-dependent STAT3 activation was observed in JAK2VF^+^ or EV cells expressing the gp130-ΔBox1 or gp130-W652A variants.

In conclusion, transduced JAK2VF^+^ cells carried more gp130 on their surface compared to control cells. This effect was absent when the Box1 motif was deleted, indicating that interaction between gp130 and JAK2VF is crucial for the observed effects. In line with these findings, IL-6 mediated activation of STAT3 was absent in cells transduced with gp130-ΔBox1. Furthermore, no IL-6/sIL-6Ra response was detected in gp130-W652A cells as already described [20].

### 2.5 Increased gp130 surface expression and IL-6 response is caused by the VF mutation in JAK2 and not total JAK2 levels

Initially, 32DEV cells were used as controls to investigate gp130-JAK2VF signaling (Figure 2, Figure 3, and Figure 4). However, published data suggests that overexpression of JAKs can lead to signaling artifacts [20]. To verify that the observed effects were due to JAK2VF activity and not caused by differences in overall JAK2 protein levels, 32D cells were transfected with JAK2-WT (32DJAK2). Western-blot analysis confirmed similar JAK2 expression in both JAK2^+^ and JAK2VF^+^ cells, compared to untransduced cells, which express quite low levels of endogenous JAK2 (Figure 5A).

**Figure 5:**
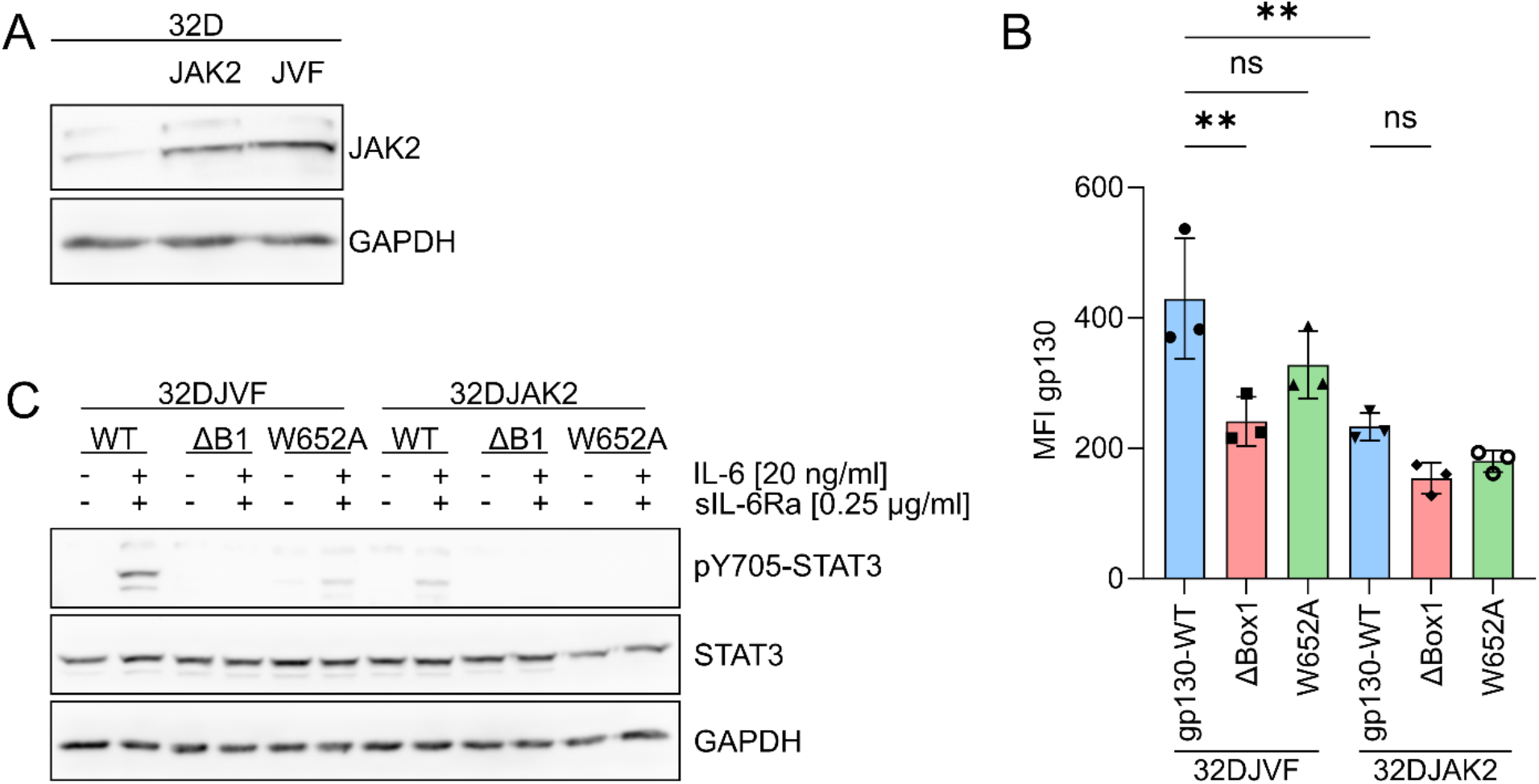
Increased gp130 surface expression and elevated IL-6-mediated STAT3 activation are not observed in 32DJAK2 cells. 32DJVF and 32DJAK2 cells were transduced with gp130-WT, gp130-ΔBox1 or gp130-W652A. **(A)** Cell lysates were prepared for SDS-PAGE and Western-blot analysis of JAK2. The detection of GAPDH served as a control. **(B)** Gp130 surface expression was analyzed via flow cytometry and the MFI for gp130 was calculated in the indicated cell lines. **(C)** Transduced 32DJVF and 32DJAK2 cells were stimulated with 20 ng/ml IL-6 and 0.25 µg/ml sIL-6Ra for 30’ at 37 °C. Cell lysates of unstimulated and stimulated cells were prepared for SDS-PAGE and Western-blot analysis of pY705-STAT3. Detection of STAT3 and GAPDH served as controls. Data was analyzed via FlowJo^TM^ v10 as well as GraphPad Prism v10. P-values were calculated via one-way ANOVA with Tukey multiple comparison test (p<0.05*, p<0.0001****).

32DJAK2 cells were then transduced with gp130-variants and flow cytometry analysis as well as IL-6 stimulation were performed. We observed significantly elevated surface expression of gp130-WT cells in JAKVF^+^ cells compared to JAK2-WT cells (Figure 5B). Furthermore, gp130-ΔBox1 surface expression was again reduced significantly in JAK2VF^+^ cells compared to gp130-WT. Next, gp130-transduced 32DJAK2 and 32DJVF cells were stimulated with IL-6/sIL-6Ra. In line with previous experiments, stronger IL-6-mediated STAT3 activation was observed in 32DJVF cells compared to 32DJAK2 cells expressing gp130-WT (Figure 5C and Figure S6). No IL-6-mediated induction of STAT3 phosphorylation was observed in cells expressing gp130-ΔBox1 or gp130-W652A.

In conclusion, increased gp130 surface expression and IL-6 response of gp130-transduced 32DJAK2VF cells is a consequence of the VF mutation and not due to JAK2 overexpression.

### 2.6 Elevated IL-6-mediated STAT3 activation in JAK2VF+ MPN patient cells compared to healthy donors

We aimed to verify increased IL-6 sensitivity in JAK2VF^+^ MPN patient cells compared to healthy donors (HD). PBMCs derived from MPN patients or HDs were stimulated with IL-6 in combination with sIL-6Ra and induction of pY705-STAT3 was analyzed via intracellular flow cytometry. The mean fluorescence intensity (MFI) of pY705-STAT3 in every sample was determined as a measure of STAT3 activation. MFI values of stimulated samples were normalized to the respective unstimulated samples (Figure 6). JAK2VF^+^ patient cells demonstrated a significantly higher fold-change in pY705-STAT3 MFI upon stimulation compared to HD samples.

**Figure 6:**
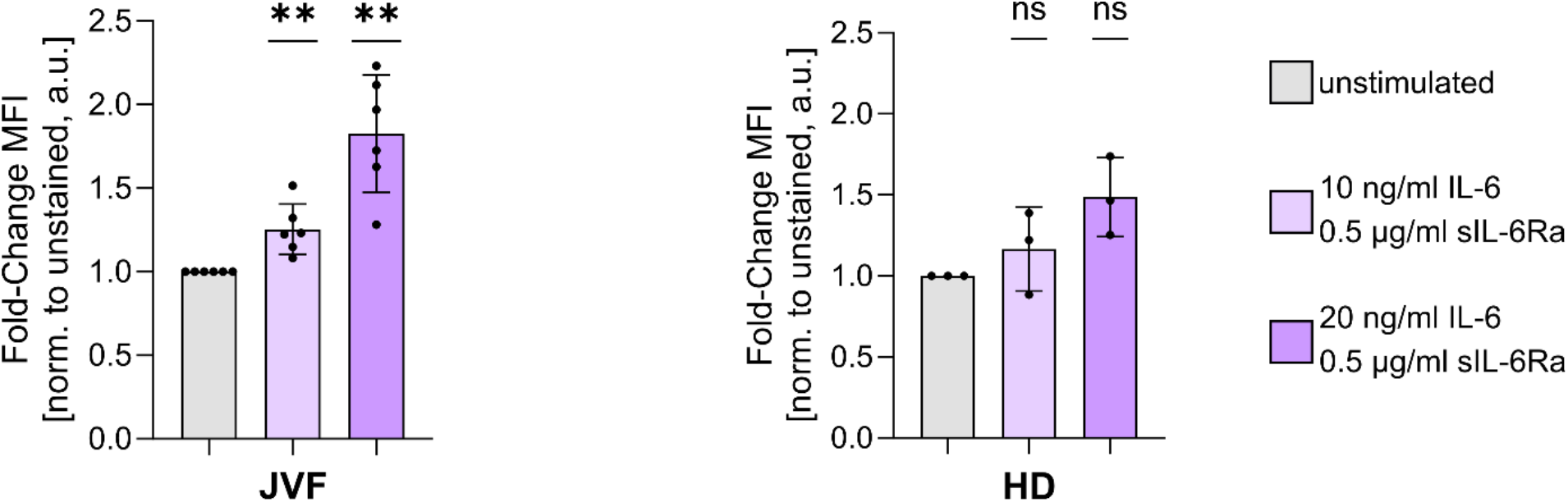
JAK2VF^+^ patient cells are more sensitive to IL-6 stimulation than HD cells. PBMCs were isolated from MPN patients or HDs and subsequently cultured for two weeks in medium containing IL-6, FLT3L, SCF and IL-3. All cells were starved in cytokine-free medium supplemented with 2% FCS for 2 h. Cells were stimulated with 10 or 20 ng/ml IL-6 in combination with 0.5 µg/ml sIL-6Ra for 30’ at 37 °C. All samples were harvested for flow-cytometry of pY705-STAT3 MFI values of pY705-STAT3 were normalized to the respective unstimulated samples to calculate the fold-change. P-values were calculated via one sample t-test with a hypothetical value of 1 (p<0.01**). Data was analyzed via FlowJo^TM^ v10 as well as GraphPad Prism v10.

In conclusion, the increased IL-6 sensitivity determined in JAK2VF transduced 32D cells is also observed in patient cells.

## 3. Discussion

Inflammatory cytokines play an essential role in the development and progression of MPNs, including PV, ET, and PMF. These diseases are characterized by chronic overproduction of pro-inflammatory cytokines such as IL-1β, TNF-α, IL-6, and others, which drive a persistent state of inflammation in the bone marrow microenvironment [30]. Elevated cytokines contribute to abnormal blood cell proliferation, bone marrow fibrosis, and clinical symptoms such as fever, fatigue, and night sweats [31]. We asked the question whether the prominent MPN driver mutation JAK2VF not only promotes cytokine production but also enhances the sensitivity of mutated cells to cytokine stimulation. We focused on IL-6 because of its central role as a pro-inflammatory cytokine in inflammation and cancer [32] and the known interaction of its signal-transducing receptor subunit gp130 with JAK2 [10].

While the interaction of gp130 with JAK1, JAK2 and also TYK2 is firmly established, their relative contribution to gp130-mediated signaling is less well defined. JAK1 appears to be most prominently involved as its deletion results in severe impairment of IL-6 signaling [9, 21]. However, involvement of JAK2 in gp130 signaling might be cell type-specific [26], potentially depending on the relative expression levels of JAK1 and JAK2. A recent study employing an artificial insect cell system reported comparable binding affinities of gp130 for JAK1 and JAK2, with lower affinity for TYK2 [25]. Further mechanistic insights into how the VF mutation renders JAK2 constitutively active have been revealed [13]: substitution of valine with phenylalanine enhances the dimer interface within the JH2 domain.

Using structural modeling, we analyzed the relative interactions of gp130 with JAK1 and JAK2, alongside other cytokine receptors that form homodimers with JAK2 (Figure 1). While most homodimeric receptors appear to preferentially recruit JAK2 over JAK1, gp130 has been reported to exhibit the opposite tendency. This is a trend that our modelling approach was able to capture. Notably, our modelling still predicts similarly stable complexes for both JAK1 and JAK2. Moreover, we examined how the increased dimer interface affinity in JAK2VF may affect the stability of receptor–JAK complexes. We modelled both the wild-type JAK2 dimer and the JAK2V617F mutant dimer. Our molecular dynamics analysis of the pseudokinase domain, which contains the mutation site and directly contributes to the dimer interface, showed that the mutation strengthens dimer stability. This is consistent with previous reports [13]. This enforced dimer persistence could explain the increased JAK2 activity, but it could also promote the formation of a more stable JAK2-receptor tetramer. In the case of gp130-JAK2VF, the augmented JAK2 dimerisation and subsequent gp130-JAK2VF tetramerization could increase the stability of the signaling complex facilitating enhanced downstream signaling upon IL-6 binding.

Indeed, the IL-6-induced STAT3 activation was more pronounced in 32DJVFgp130 compared to 32DEVgp130 cells (Figure 2) suggesting an increased IL-6 sensitivity of JAK2VF^+^ cells. Wilmes and colleagues found that JAK2VF induced ligand-independent dimerization of homodimeric cytokine receptor complexes like TPOR and EPOR [33]. More specifically, the JAK2VF mutation induces a more stable dimer interface between receptor-associated JAKs which is reminiscent of the ligand-induced conformational change [13]. Similar to the EPOR or TPOR, JAK2VF might mimic ligand independent dimerization of gp130 chains as suggested by our initial modeling studies (Figure 1). Consequently, a higher frequency of pre-formed gp130 dimers in 32DJVFgp130 cells might facilitate an increased sensitivity towards IL-6 and lead to increased STAT3 activation.

As JAK1 is generally considered to be the canonical kinase bound to gp130 and activated upon IL-6 stimulation [9, 21], JAK1-KO 32D cells were generated via CRISPR/Cas9 genome editing to prevent competition between gp130-JAK1 and gp130-JAK2VF signaling. (Figure 3 and Figure S2). IL-6-mediated STAT3 activation in JAK1-KO 32DJVFgp130_CRISPR_ cells suggested that the elevated activation of STAT3 is independent of JAK1 (Figure 3A). Furthermore, the IL-6-induced activation of STAT3 in 32DEVgp130_CRISPR_ cells confirms that JAK2 can generally substitute for JAK1 in IL-6 signaling.

To substantiate the findings made with JAK1-KO cells, cells were treated with the JAK2 specific inhibitor BMS prior to IL-6 stimulation. BMS is an ATP competitive inhibitor highly selective for JAK2 (IC_50_= 1.1 nM) [34]. IL-6-mediated activation of STAT3 was abrogated upon BMS treatment in both JAK1-WT and JAK1-KO cells suggesting that JAK2VF activity mediates the increased induction of pY705-STAT3 (Figure 3B). Despite the high selectivity of BMS for JAK2, off-target effects might always be present, especially since higher BMS concentrations (1 µM) were necessary for JAK2 inhibition *in vitro* than the expected from the suggested IC_50_ of 1.1 nM. However, possible JAK1 off-target effects were not relevant in JAK1-KO cells.

Gp130 surface expression was significantly increased in 32DJVFgp130 compared to 32DEVgp130 or 32DJAK2gp130 cells (Figure 4, Figure 5, and Figure S4). The influence of JAKs on cytokine receptor transport and processing is well studied. EPOR processing in the endoplasmic reticulum/Golgi pathway is positively modulated by JAK2 leading to increased surface presentation of EPOR [35]. Similarly, TPOR surface expression is mediated by JAK2 and TYK2, which both stabilize the mature TPOR [36]. Furthermore, JAK2VF has been reported to deregulate receptor surface expression. Platelets from MPN patients exhibit decreased surface expression of the TPOR receptor [37]. Mechanistically, JAK2VF directly reduced TPOR expression via increased ubiquitination and reduced recycling as well as reduced maturation of the receptor [29]. Our findings in 32DJVFgp130 cells further support the influence of MPN oncogenes on receptor surface expression.

Gp130 is almost ubiquitously expressed. 32D cells belong to the rare cell lines that do not express gp130 endogenously. Therefore, these cells are perfectly well suited for the analysis of gp130 mutants because no interference with gp130 wild type does occur as in most other cell lines. We transduced 32D cells with gp130 variants mutated in the critical JAK interaction motif [20]. Surface expression of gp130-ΔBox1 was significantly reduced compared to gp130-WT in transduced 32DJVF cells and no IL-6 mediated STAT3 activation was detected in 32DJVFgp130-ΔBox1cells (Figure 4 and Figure 5). These findings strongly suggest that increased surface expression of gp130 depends on the interaction with JAK2VF.

After demonstrating enhanced gp130 surface expression and IL-6 sensitivity in JAK2VF^+^ 32D cell lines, we next investigated the IL-6 response in cells derived from JAK2VF^+^ MPN patients. IL-6 stimulation of JAK2VF^+^ MPN patient cells resulted in a stronger STAT3 activation compared to IL-6 stimulation of HD cells (Figure 6) underlining the clinical relevance of our previous findings in 32D cells. Recently, it was shown that cells carrying CALRdel52, another MPN driver mutation, are more sensitive to IL-6 signaling as well [7].

Collectively, our findings expand the current understanding of the functional impact of the JAK2VF mutation on cytokine receptors. Our *in silico* modeling revealed that the JAK2VF mutation exhibits an increased tendency to dimerize, potentially enhancing its affinity to gp130 and promoting the formation of gp130-JAK2VF tetramers. Cells expressing gp130-JAK2VF showed higher gp130 surface expression and increased IL-6 signaling compared to gp130-JAK2 expressing cells. In the context of MPNs, augmented IL-6 signaling in myeloid cells could amplify inflammatory responses and exacerbate disease symptoms, including progression to fibrosis. These insights suggest that aberrant gp130-JAK2VF interactions and enhanced IL-6 signaling may represent novel therapeutic vulnerabilities in MPNs, warranting further exploration of targeted strategies to mitigate inflammation-driven disease progression.

## 4. Material and methods

### 4.1 Modeling and scoring of kinase-receptor interactions

Full sequences of human kinases (JAK1/P23458, JAK2/O60674) were retrieved using the UniProt API [38]. Domain boundaries were defined to extract the FERM and SH2 domains based on structural data including pLDDT (predicted local distance difference test) and PAE (predicted align error) scores for the kinases from the AlphaFold protein structure database [39, 40]: Jak1/1-559, JAK2/1-514. Earlier defined intracellular parts of receptors containing conserved Box1 and Box2 were used [12].

*Ab initio* structure and interaction prediction of all kinase-receptor pairs of interest was performed using the deep learning algorithm AlphaFold Multimer (AFM) [40] in a high-throughput batch pipeline based on the ColabFold notebook v1.5.5 [41]. The modelling was performed without homology templates, using MMseq2 for multiple sequence alignment. Four predictions (seeds = 4) per AlphaFold model were produced resulting in 20 structural models per dimer pair. The predicted models were ranked according to the ipTM (interface predict template model) score. To calculate the interface PAE (iPAE) for each dimer model, a custom Python code was used to extract CA atom coordinates from both chains to identify pairs of residues within 10 Å of each other. These pairs were then clustered into interfaces based on spatial continuity. For each identified interface, the average iPAE was calculated by combining the PAE values of all involved residue pairs.

FoldX5 [42] was used for (a) structural model refinement and (b) model quality analysis based on the absence of backbone collisions. Ten consecutive rounds of side-chain structural relaxation were performed to minimise the energy of the kinase-receptor dimer models. Models with backbone clashes were classed as failed models as no useful further analysis could be performed. After refinement, FoldX5 force field was used to calculate interaction energies (at an ionic strength of 150 mM).

Furthermore, relaxed models were used to assess predicted dimer interaction with the machine learning model VoroIF-GNN (Voronoi tessellation-derived protein– protein interface assessment using a graph neural network) [22].

All python scripts are available upon reasonable request and after publication from GitHub repository (https://github.com/Clusterbiology).

The predicted AFM models were visualised using ChimeraX (v1.7.1) [43]

### 4.2 Molecular dynamics simulation

Molecular dynamics simulation was used to analyse the dynamics of the JAK2 PK-PK dimerization interface over time. The best ranked AlphaFold model of the dimer was used. The system was immersed in TIP3P water in a cubic box with 1.0 nm padding and neutralised with 150 mM NaCl. The MD simulation was performed with GROMACS2023.2 [44] using the AMBER99SB-ILDN force field [45]. The MD system was energy minimised and equilibration simulations were performed with periodic boundary conditions. The temperature was equilibrated using an *NVT* ensemble for 1 ns followed by an *NPT* ensemble for 1 ns. Finally, the production MD simulations were performed for 100 ns at 300 K and 1 atm.

### 4.3 Reagents

Dulbecco’s modified eagle medium (GlutaMAX^TM^, DMEM) and RPMI1640 medium were purchased from Gibco^TM^ (Invitrogen). Fetal calf serum (FSC) was purchased from Capricorn Scientific and heat-inactivated before use (LOT:CP32-4515). Penicilin/Streptomycin (P/S) was purchased from Sigma. Puromycin and Blasticidin were sourced from InvivoGen. IL-3 for maintenance culture was derived from conditioned medium of X63Ag8653-BPV-mIL-3 cells. Recombinant murine (rm) and recombinant human (rh) cytokines were all purchased from ImmunoTools. TransIT-LT1 transfection reagent was bought at MirusBio and polybrene was purchased from Sigma-Aldrich. The Retro-X^TM^ concentrator was purchased from TakaraBio. The JAK2-specific inhibitor BMS-911543 was sourced from MedChemExpress (*REF for inhibitor*). IDTE buffer, Alt-R^TM^ Cas9 electroporation enhancer, Alt-R^TM^ tracrRNA-Atto550, gRNAs, Alt-R^TM^ S.p. Cas9 Nuclease V3 and nuclease free duplex buffer were bought at Integrated DNA technologies.

### 4.4 Cell culture

Platinum-E (Plat-E, RRID: CVCL_B488) cells were maintained under standard culture conditions in DMEM supplemented with 10% FCS and 1% P/S together with puromycin (1 µg/ml) and blasticidin (15 µg/ml). 32D cells (RRID: CVCL_0118) were cultured in RPMI1640 supplemented with 10% heat-inactivated FCS, 1% P/S and IL-3 (self-made). Cells transduced with puromycin encoding transfer vectors were additionally cultured in the presence of puromycin (1 µg/ml). For all stimulation experiments, 32D cells were cultured with a defined concentration of rmIL-3 (2 ng/ml). All cell lines were regularly tested for mycoplasm contamination. All experiments were performed with mycoplasm-free cells.

Cells derived from MPN patients or HDs were collected at the Department of Hematology, Oncology, Hemostaseology and Stem Cell Transplantation and the Institute of Transfusion Medicine (Faculty of Medicine, University Hospital, RWTH Aachen University) and were obtained after informed and written consent, as approved by the local ethics committee (EK127/12, EK099/14). Stimulation experiments were carried out with MPN patient cells or HDs cultured in RPMI1640 supplied with rhSCF (100 ng/ml), rhIL-6 (25 ng/ml), rhIL-3 (30 ng/ml) and rhFlt3L (25 ng/ml) for two weeks prior to analysis.

### 4.5 Generation of 32D cells expressing EV or JAK2VF and gp130 variants

32DEV and 32DJVF cells as well as the MSCV-mJAK2-IRES-GFP vector (Addgene, #20672) were a kind gift from our collaboration partners at the Clinic for Hematology, Oncology, Hemostaseology and Stem-cell Transplantation (University Hospital, RWTH Aachen University). 32DJAK2 cells were generated via γ-retroviral transduction. To this end, Platinum-E cells were transfected with the MSCV-mJAK2-IRES-GFP vector using TransIT-LT1 according to the manufacturer’s instructions. Three days after transfection, virus containing supernatant was harvested from Platinum-E cells using the Retro-X^TM^ concentrator according to the manufacturer’s instructions. Concentrated virus was added to target cells in combination with polybrene (8 µg/ml) followed by incubation under standard conditions over night. A second transduction step was performed for 6 hours on the next day.

The coding sequences of wildtype (WT) human gp130 (hgp130) as well as the ΔBox1 and W652A variants were derived from the work of Haan and colleagues and subsequently cloned into the MSCV-puro vector (Addgene, #68469) (REF). Generation of 32DEV, 32DJAK2 and 32DJVF cells encoding the various gp130-variants was performed via retroviral transduction as described above.

### 4.6 Flow cytometry and fluorescence-activated-cell-sorting (FACS)

For flow cytometry and FACS, cells were harvested and washed with FACS-buffer (PBS, 5% FCS, 0.1% NaN_3_). Staining of target cells was performed for 25 minutes at 4 °C in the dark. To analyze gp130 surface expression, target cells were stained with a gp130-specific antibodies which were either conjugated to PE (R&D, FAB228P-025) or unlabeled (Diaclone), Unlabeled antibodies were used in combination with an anti-mouse-Alexa647 antibody (Invitrogen, A212236). For intracellular flow cytometry, samples were fixed with 4% paraformaldehyde and permeabilized with ice-cold methanol. STAT3 activation was analyzed by staining cells with a phosphor-STAT3-Y705 antibody (Cell Signaling Technology, 9145S) in combination with an anti-rabbit-Alexa488 antibody (Invitrogen, A11034). Flow cytometry was performed with the FACSCanto^TM^ II (BD Biosciences).

FACS was performed at the FACSAria^TM^ Fusion (BD Biosciences) under sterile conditions at 37 °C. Experimental assistance during FACS was provided by the Flow Cytometry Facility (IZKF, RWTH Aachen University Hospital). Cells were sorted into tubes (bulk-sorting) or 96-well TC-dishes (single-cell sorting) for further expansion.

### 4.7 CRISPR/Cas9 genome editing

Predesigned crRNAs were sourced via the free-to-use web-tool by IDT^TM^ (https://eu.idtdna.com/site/order/designtool/index/CRISPR_SEQUENCE). The spe-cies was set to “Mus musculus” and “gene symbol” was chosen as the input format (Table 1). To assemble the gRNA, an equimolar mix of crRNA and tracrRNA was incubated at 95 °C for precisely 5 minutes. Next, gRNA and Cas9 were incubated together for 30 minutes at room temperature to assemble the ribonucleoprotein (RNP)-complex. Electroporation (Nucleofection®, Lonza) of the RNP-complex was performed with the SF Cell Line 4D-NucelofectorTM X Kit (Lonza, #V4XC-2032) according to the manufacturer’s instructions. Nucleofection® was performed at the 4D-Nucleofactor^TM^ (Lonza) with a cell line specific program recommended by the manufacturer. FACS for Atto550^+^ cells was performed to generate single-cell populations (Figure S2A). Single-cell clones were expanded, and successful genome-editing was verified using PCR, Sanger-sequencing of gDNA as well as SDS-PAGE and Western-blot analysis.

**Table 1:**
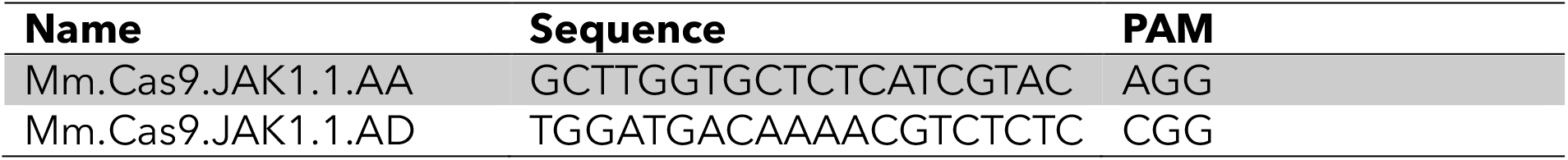
List of crRNAs used for the generation of CRISPR/Cas9-mediate gene-editing.

### 4.8 DNA isolation, PCR and Sanger-sequencing

DNA was isolated from cells using the QIAmp DNA Mini Kit (Qiagen, #56304) according to the manufacturer’s instructions. PCR was performed via a standard Taq-polymerase protocol using JAK1 specific primers (5’-TTATATCCGTGGTCCTCAGG-3’, 5’-CAGCACATTCCCTTGATGAC-3’). Sanger-sequencing was performed by Eurofins Genomics. Samples were prepared according to the company’s guidelines via the Mix2Seq kit. Sequencing results were aligned to a reference sequence using SnapGene software (https://www.snapgene.com/). For gDNA analysis of clones generated during CRISPR/Cas9-mediated genome editing, sequencing results were analyzed using ICE analysis v3.0 (Synthego, https://ice.synthego.com/).

### 4.9 Cell lysis and immunoblotting

Cell lysates were prepared via standard RIPA-lysis for SDS-PAGE and Western-blotting. Membranes were probed with antibodies directed against phospho-Stat3 (Tyr705, Cell Signaling Technologies, 9131S), Stat3 (Cell Signaling Technologies 9139S), phospho-Jak2 (Tyr1007/1008, Cell Signaling Technologies, 3771S), Jak2 (Cell Signaling Technologies, 3230S), Jak1 (Cell Signaling Technologies, 29261S) and GAPDH (SantaCruz, sc-32233). Anti-mouse-HRP (Agilent/Dako, P0477) or anti-rabbit-HRP (Agilent/Dako, P0488) were used as secondary antibodies. Immunodetection was performed with pCA-ECL + H_2_O_2_ (0.1 M Tris/HCl pH 8.8, 2.5 mM Luminol, 0.2 mM p-coumaric acid, 10 mM H_2_O_2_) at the “LAS4000 Mini” (Fuji Film).

### 4.10 Data analysis

Statistical analysis was performed via Graphpad Prism 10. Data were analyzed using a one-way ANOVA with Tukey’s multiple comparison test or a one-sample t-test with a hypothetical value of 1. Results were considered significant when a *P*-value of

< 0.05 was reached. Flow cytometry data was analyzed using FlowJo v10.

## Supporting information

Supplement Schurse etal

## 5. Author contributions

H.S.: investigation, visualization, methodology, data analysis, manuscript–original draft, writing–review, and editing. M.A.S.T.: resources, methodology, writing-review, and editing. A.K.: methodology, writing-review, and editing. S.K. resources, writing-review, and editing. M.C.: resources, funding acquisition, writing-review, and editing. S.D.: investigation, visualization, methodology, funding acquisition, manuscript– original draft, writing–review, and editing. G.M.-N.: conceptualization, supervision, resources, funding acquisition, project administration, formal analysis, writing-review, and editing.

## 6. Acknowledgements

This study was funded by grants to G.M.-N. from the German Research Foundation (Deutsche Forschungsgemeinschaft, DFG, MU 1331/4-1 and MU 1331/4-2; project number 417911533), to S.D. from the German Federal Ministry of Education and Research (01KD24235) and to S.K. by funds from the German Research Foundation as part of the Clinical Research Unit CRU 344 (KO2155/7-1; project number 428858786). We would like to thank Hildegard Schmitz-van de Leur for plasmid cloning. The authors would also like to thank all the patients who kindly donated samples.

## Notes

### Competing Interest Statement

The authors have declared no competing interest.

