## Supplement Schurse etal for "The V617F mutation in JAK2 renders myeloid cells more sensitive to IL-6-mediated gp130 signaling"

### Supplemental Figures

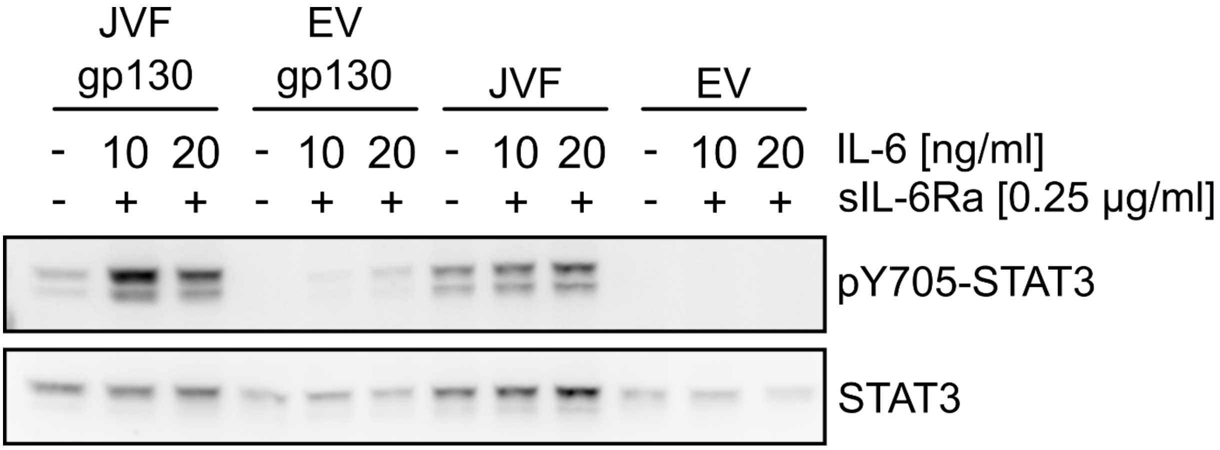


**Figure S1: Increased IL-6 response in 32D cells upon expression of JAK2VF.** Replicate of Western-blot shown in Figure 2. Transduced and untransduced cells were stimulated (10 or 20 ng/ml IL-6, 0.25 µg/ml sIL-6-Ra, 30' at 37 °C) and cell lysates were prepared for SDS-PAGE and Western-blot analysis. The membrane was probed with a pY705-STAT3 antibody and detection of STAT3 or served as a control.

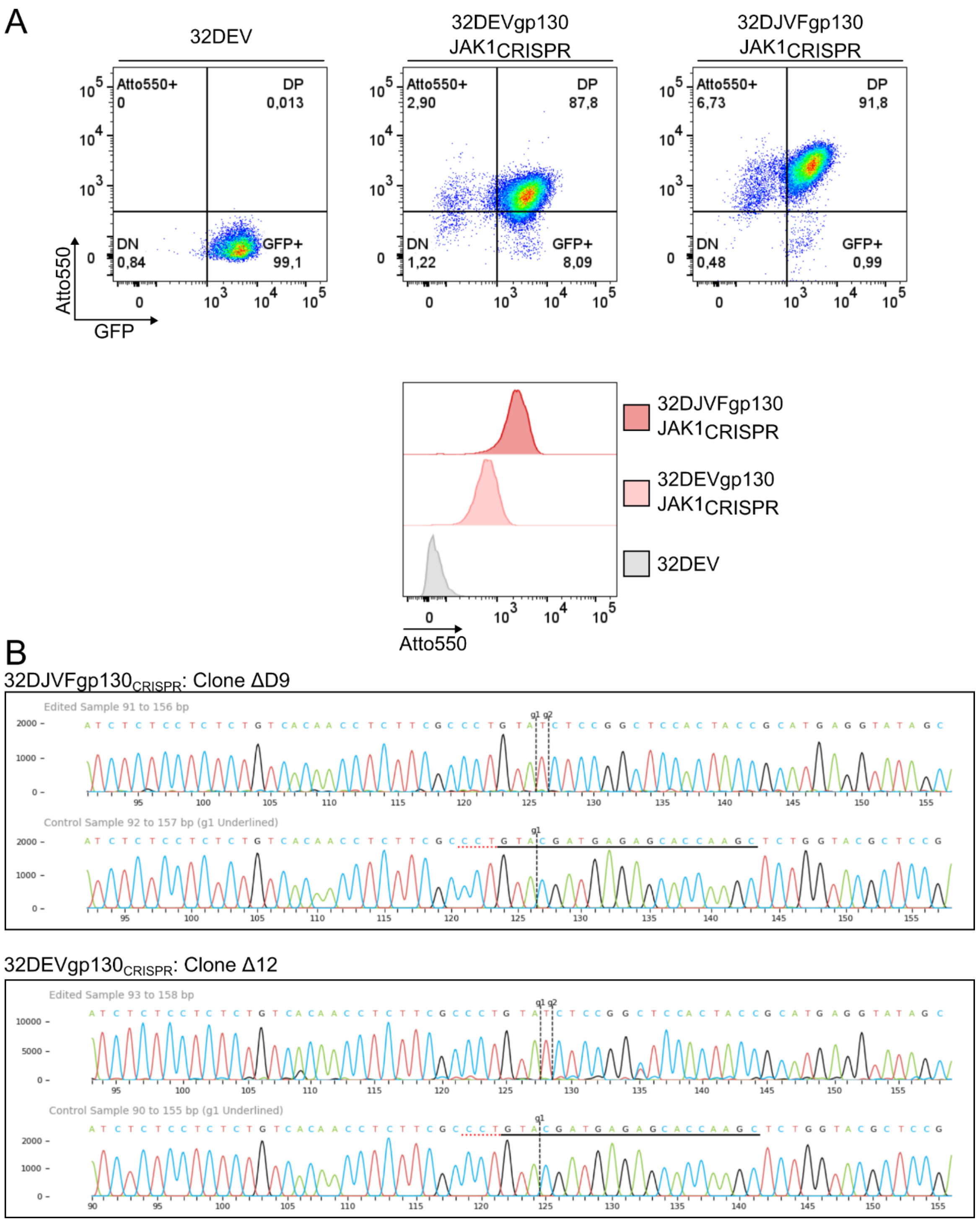


**Figure S2: CRISPR-Cas9-mediated knockout of JAK1 in 32DEVgp130 and 32DJVFgp130 cells.** Atto550-labelled gRNA and Cas9 were assembled into an RNP-complex, which was subsequently nucleofected into 32DEVgp130 and 32DJVFgp130 cells. **(A)** Atto550$^{+}$GFP$^{+}$ clones from nucleofected cells were enriched via single-cell FACS. 32DEV cells were used as a control. **(B)** gDNA was isolated from single-cell clones a region around the gRNA-target site was PCR-amplified. Sanger sequencing was subsequently performed, and sequences were analyzed using ICE. 2019. v3.0. Synthego [11/Sep/2024]. Representative reads are shown for one clone of 32DJVFgp130$_{CRISPR}$ (ΔD9, upper panel) and 32DEVgp130$_{CRISPR}$ cells (Δ12, lower panel).

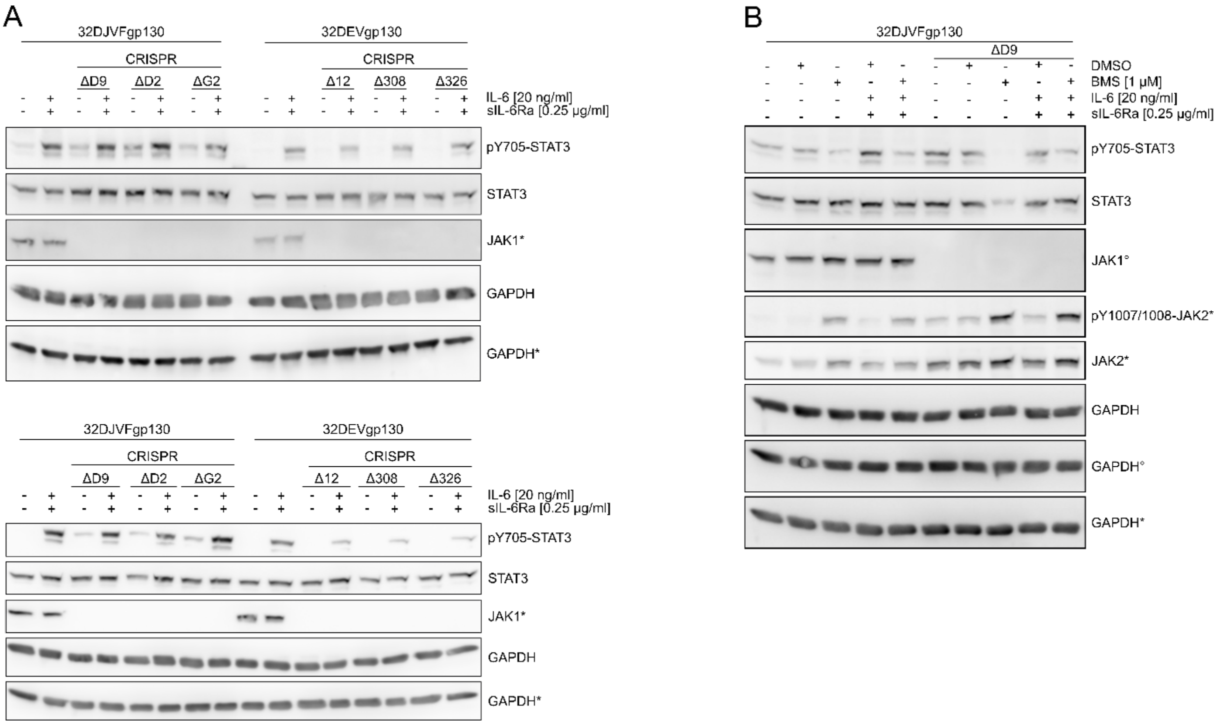


**Figure S3: Increased IL-6-mediated STAT3 activation in 32DJVFgp130 cells is independent of JAK1 but sensitive to inhibition of JAK2.** Replicates of Western-blots shown in Figure 3. **(A)** Cell lysates were prepared from unstimulated and stimulated (20 ng/ml IL-6 and 0.25 µg/ml sIL-6Ra for 30′ at 37 °C) cells. SDS-PAGE and Western-blot analysis were performed for p705-STAT3 and JAK1. Detection of STAT3 and GAPDH served as controls. **(B)** Cells were treated with BMS for 16h prior to stimulation. Cell lysates were prepared for SDS-PAGE and Western-blot analysis of pY705-STAT3, pY1007/1008-JAK2 and JAK2. Detection of JAK2, STAT3 and GAPDH served as controls. Different immunoblots were used to detect STAT3 and JAKs. Corresponding loading controls are marked by * or °.

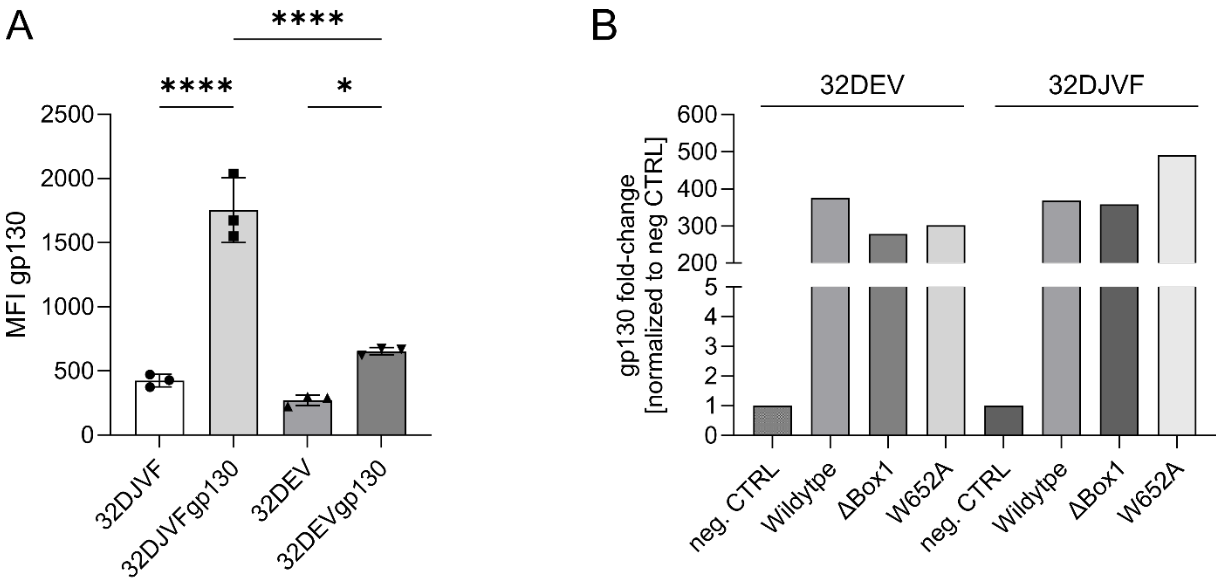


**Figure S4: Flow cytometry and qPCR analysis of gp130 expression in transduced 32D cells.** 32DJVF and 32DEV cells were transduced with gp130-WT, gp130-ΔBox1 or gp130-W652A. **(A)** Surface expression of gp130 was analyzed via flow cytometry in untransduced and transduced cells with subsequent calculation of the MFI for gp130. **(B)** RNA was isolated from untransduced and transduced cells and converted into cDNA for RT-qPCR. The fold-change was calculated using normalization to GAPDH and the ΔΔCT method with untransduced cells as the reference. Data was analyzed via FlowJo™ v10 and GraphPad Prism v10. P-values were calculated via one-way ANOVA with Tukey multiple comparison test ($p<0.05$*, $p<0.0001$****).

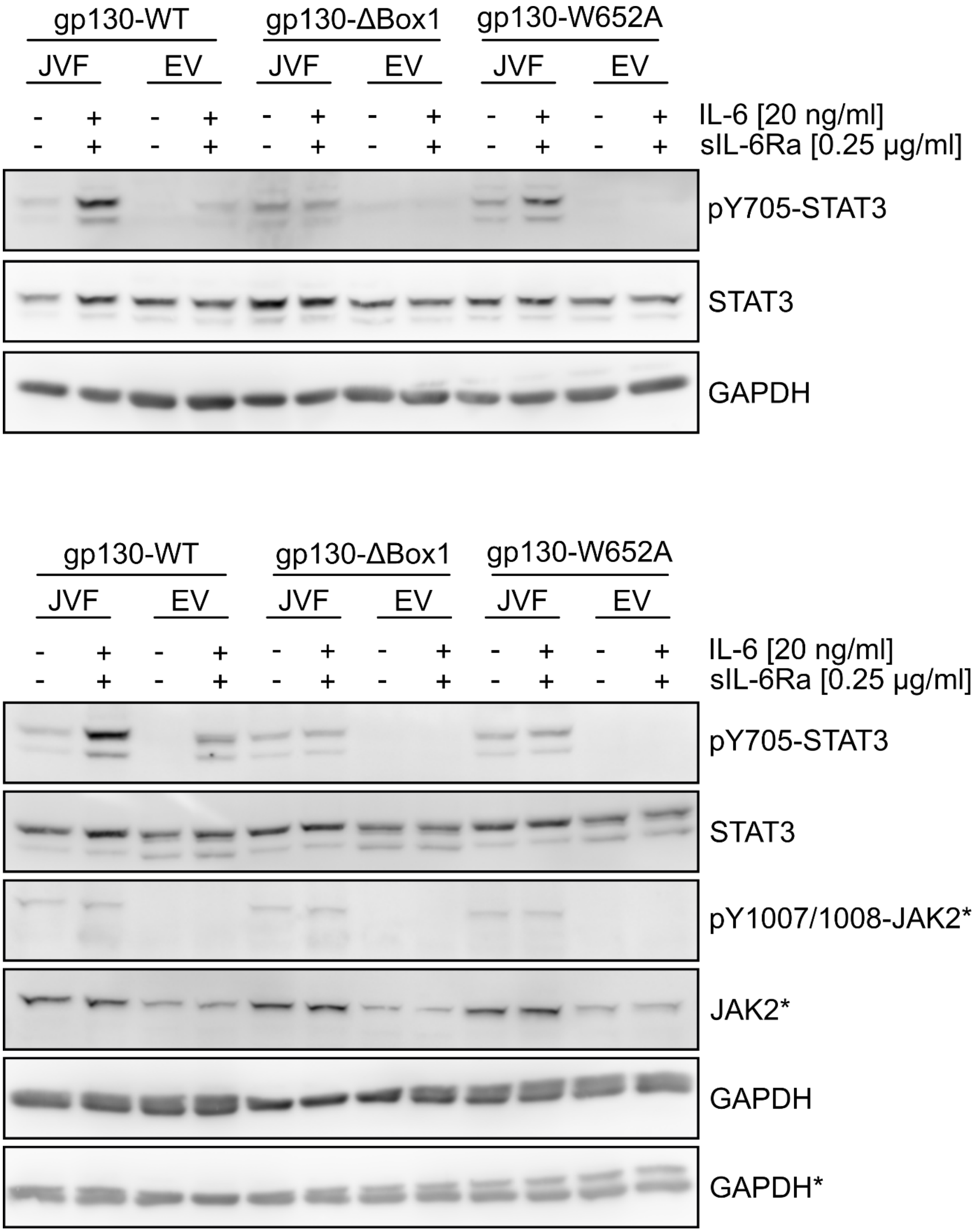


**Figure S5: The JAK interaction motif in gp130 is necessary for elevated IL-6 response.** Replicates of the Western-blot shown in Figure 4. Indicated cell lines were stimulated with 20 ng/ml IL-6 and 0.25 µg/ml sIL-6Ra for 30′ at 37 °C. Cell lysates of unstimulated and stimulated cells were prepared for SDS-PAGE and Western-blot analysis of pY705-STAT3 and pY1007/1008-JAK2. Detection of STAT3, JAK2 and GAPDH served as controls. Different immunoblots were used to detect STAT3 and JAK2. Corresponding loading controls are marked by *.

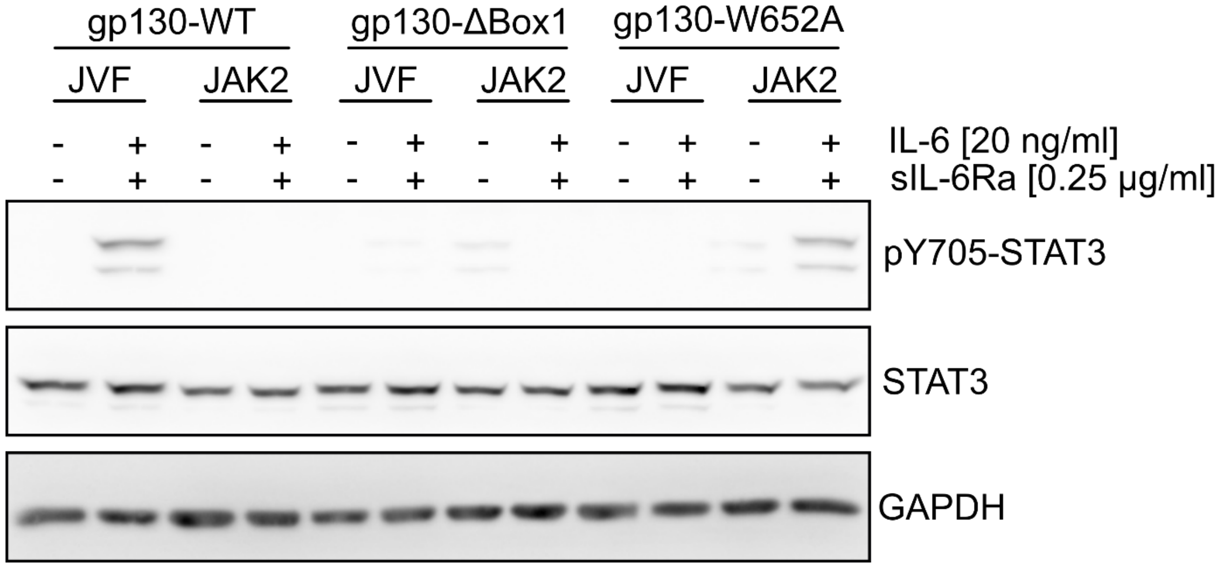


**Figure S6: No increase of IL-6-mediated STAT3 activation observed in 32DJAK2 cells.** Replicate of the Western-blot shown in Figure 5. Transduced 32DJVF and 32DJAK2 cells were stimulated with 20 ng/ml IL-6 and 0.25 µg/ml sIL-6Ra for 30′ at 37 °C. Cell lysates of unstimulated and stimulated cells were prepared for SDS-PAGE and Western-blot analysis of pY705-STAT3. Detection of STAT3 and GAPDH served as controls.
